# Calcium-phosphate bridge is a novel phosphorylation switch that stabilises protein-complexes during HIV assembly

**DOI:** 10.64898/2026.02.25.708077

**Authors:** Erwan Brémaud, Biswa Prasanna Mishra, Abdulhakim Bake, Veronika Masic, Thibaut Flipo, Arun Everest-Dass, Lauren Hartley-Tassell, Paul R Gorry, Thomas Ve, Belinda L Spillings, Johnson Mak

## Abstract

Calcium (Ca^2+^) and phosphate (PO_4_^3-^) are fundamental-element and -chemical group in biology. Specifically, the chemistry of both Ca^2+^ signalling and phosphorylation switch are independent mechanisms regulating a broad spectrum of biological processes. It is, however, not appreciated that a normal function of phospho-mimic amino acids (aspartate/glutamate) is to interact with Ca^2+^ at the atomic level. Here, we leveraged HIV-Ca^2+^ biology in primary cells to describe an unknown layer of regulatory processes via Ca^2+^-phosphate (PO_4_^3-^) bridge to support protein complex formation. We identified novel HIV phosphorylation sites overlapping Ca^2+^ binding domains through phospho-proteomics. Integrating primary cells, molecular virology, structural biology, biophysical and ultrastructural analyses, we presented multiple examples of Ca^2+^-PO_4_^3-^ bridges that support HIV assembly and function. These include Ca^2+^-PO_4_^3-^ bridges: (i) stabilising Pr55^Gag^-Pr160^GagPol^ complex for virus function; (ii) mediating p6^Pol^ dimerization to support virion maturation; and (iii) modulating viral complex formation to package both viral enzymatic- and cellular-proteins. As the convergent enrichment of these signatured calcium-phosphorylation domains occurs across a wide range of viral and cellular proteins, we propose Ca^2+^-PO_4_^3-^ bridge to be a general principle for Ca^2+^-coordinated phosphorylation switch to regulate biological processes.

## INTRODUCTION

Foundation element calcium (Ca^2+^) and chemical group phosphate (PO_4_^3-^) are amongst the best-known examples in chemical biology. Ca^2+^ signalling ^1^ and phosphorylation switch ^2^ are two separate branches of cell biology regulatory frameworks that are critical across a broad spectrum of biological processes. The discovery of Ca^2+^ sparks to regulate intracellular homeostasis ^3^ highlights the dynamic nature of Ca^2+^ signals in managing diverse cellular events. On the other hand, the only recognised mechanism for phosphorylation switch is through *direct* phosphate (PO_4_^3-^)-protein interaction ^2,4^. While aspartate and glutamate are often used as phospho-mimic amino acids to dissect the biology of phosphorylation switch ^5,6^, it is less known that one of the natural functions of these phospho-mimics amino acids is to interact with divalent calcium (Ca^2+^) ion to execute biological events ^7^. Using human immunodeficiency virus (HIV) assembly and its interaction with Ca^2+^ in primary cells ^8^ as case study, here, we show how Ca^2+^ signalling and phosphorylation switch work cooperatively to regulate biological processes, specifically by forming a previously unknown Ca^2+^-PO_4_^3-^ bridge to stabilise protein complexes and perform biological function. Based on published literature, we discuss how Ca^2+^-PO_4_^3-^ bridge could be a generic mechanism across biology.

Interactions of the viral structural precursor Pr55^Gag^ with other viral and/or cellular proteins are critical for HIV assembly and release. We have recently shown that HIV piggybacks onto host cell Ca^2+^ sparks ^3^ to facilitate HIV synapse formation ^8^ (a key mechanism to support cell-cell stealth transmission of HIV within lymphoid tissue ^9–11^). HIV assembly protein Pr55^Gag^ leverages the local surge of Ca^2+^ (hundreds of μM) from calcium release units (CRUs) to promote Pr55^Gag^-oligomerization ^8^, thereby permitting progressive assembly and migration of HIV complexes to the uropods of T-lymphocytes for particle release ^8^. Structural dissection on the details of Ca^2+^-p6^Gag^ interaction at an atomic level showed that *in vitro* Ca^2+^ binding occurs near putative phosphorylation sites. Bioinformatic analyses revealed a convergent enrichment of phosphorylation- and Ca^2+^ binding-capable amino acids surrounding late domains of endosomal sorting complex required for transport (ESCRT) dependent viral genomes. HIV particle phospho-proteomic analyses confirmed the phosphorylation status across several p6^Gag^ and p6^Pol^ amino acids as novel HIV phosphorylation sites. Biophysical studies illustrated that both recombinant HIV proteins and synthetic HIV peptides utilised Ca^2+^-PO_4_^3-^ bridge to stabilise viral protein complexes *in vitro*. Molecular virology and ultrastructural analyses have identified multiple examples of Ca^2+^-PO_4_^3-^ bridge interactions that were important for the assembly and function of HIV. In addition to describing a novel phosphorylation switch mechanism by which calcium coordinates complex formation during HIV assembly, our work highlights an unprecedented pathway to delineate the underlined mechanism in phosphorylation switch biology.

## RESULTS

### Phosphorylation capable amino acids are putative calcium-binding sites

HIV particle formation requires a precise ratio of viral precursor -structural protein (Pr55^Gag^) and -enzymatic protein (Pr160^GagPol^) expression and interaction (***Fig.1A***) ^12^. We have previously identified Ca^2+^ binding amino acids (aspartate and glutamate) near HIV ESCRT binding motifs of p6^Gag^ to stabilise HIV complexes during virological synapses formation ^8^. To better define the Ca^2+^ binding properties of HIV p6^Gag^, we performed Ca^2+^-titrations by heteronuclear single quantum coherence (HSQC) nuclear magnetic resonance (NMR) spectroscopy using ^15^N-isotope L/I/F labelled p6^Gag^ peptide (***Fig.1B***). Conditions were kept identical to those used to solve the only known p6^Gag^ NMR structure (52 amino acids peptide)^13^ (PDB: 2C55). Conditions allowed p6^Gag^ amino acids assignment to HSQC peaks, tracked upon titration with increasing Ca^2+^ concentrations. The Ca^2+^ concentration-dependent chemical shift perturbations were observed predominantly for residues near the C-terminus of p6^Gag^ (from I479 to F493), with p6^Gag^-L489, -L492, and -F493 displaying the strongest shifts (***Fig.1B***). p6^Gag^ -F463, -F465 and -L483 had smaller chemical shift changes, whereas p6^Gag^-I479 and -L486 exhibited chemical shift changes mostly at high Ca^2+^ concentrations (***Fig.1B***). Residues with the strongest chemical shift perturbations were highlighted on existing NMR solved p6^Gag^ structure (PDB: 2C55) and all flanked S491, the only residue susceptible to achieve direct Ca^2+^ interaction and to explain measured shifts. AlphaFold 3 ^14^ prediction was consistent with experimental observations; Ca^2+^ was confidently assigned proximal p6^Gag^-S491 (***Fig.1C, S1***), which further emphasized Ca^2+^ binding in p6^Gag^ C-terminus region (***Fig.1B***). These data suggested that, in addition to amino acids D and E of p6^Gag8^, other p6^Gag^ amino acids could be involved in Ca^2+^ binding.

Ca^2+^ binding can be achieved via coordinating oxygen from: (i) negative-charged side chain amino acids (i.e. D & E); and (ii) polar uncharged side chain amino acids (such as N & Q) ^7^. Bioinformatic analyses across genomes of diverse ESCRT-dependent enveloped viruses ^15^ have revealed a putative evolutionary convergent enrichment of Ca^2+^ binding amino acids (D, E, N, & Q) surrounding ESCRT binding motifs (***Fig.1D***). Furthermore, phosphorylation capable amino acids (i.e. S, T, & Y) were often found being part of as well as flanking the ESCRT binding motifs (***Fig.1D***). Atomic structural comparison between *‘Ca^2+^ binding amino acids (D, E, N, & Q)’* and *‘phosphorylated amino acids (pS, pT, & pY)’* (***Fig.1E***) suggested phosphorylated amino acids could act as bidentate ligands (as seen with amino acids-D & -E) to support Ca^2+^-coordination, thereby stabilising protein complex formation ^8^. By performing phospho-proteomic analyses on tens-of-litres-equivalent of purified HIV particles, we have identified nine novel phosphorylation sites both in HIV-p6^Gag^ (-S451, -S462, -Y484, and -S498) and -p6^Pol^ (-S449, S450, -T453, -S457, and -T459) ^16^. The detected phosphorylated amino acids included previously described (or suspected) HIV-p6^Gag^ phosphorylation sites (p6^Gag^-T456, - S488, -S491, and -S499) (***Fig.1F, S2***) ^17–21^. As one would expect from any transient phosphorylation switch mechanism, none of these amino acids was 100% phosphorylated in virion particles. However, differences in prevailing sites across strains (***Fig. 1F***) and co-phosphorylation events (***Fig.1F, S2***) illustrate the anticipated transient nature of phosphorylation events in the regulation of biological processes.

**Figure 1:**
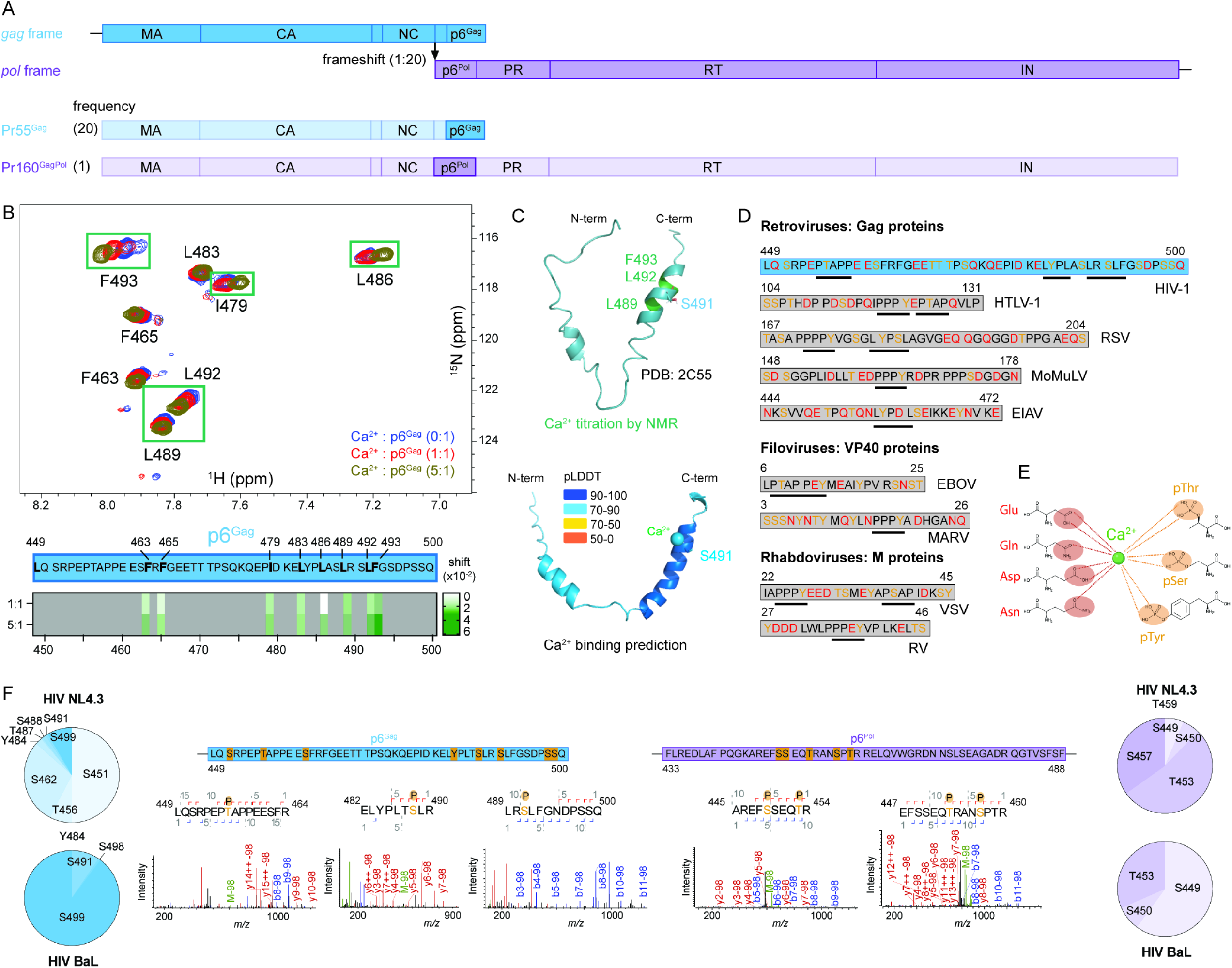
Potential contribution of phosphorylation prone p6^Gag^ amino acids to Ca^2+^-binding near ESCRT binding motifs. (A) Frameshifting in HIV genome controls expression frequencies (20:1) of Pr55^Gag^ and Pr160^GagPol^. (B) HSQC spectrum of p6^Gag^ selectively labelled for I/L/F (residues in bold in peptide sequence) showing residues undergoing chemical shift perturbation with increasing concentrations of Ca^2+^. Peaks were assigned based on published p6^Gag^ NMR structure ^13^. Ca^2+^ binding characterisation using chemical shift perturbation ^22^. Intensities of peaks shift were quantified between 1:1 or 5:1 ratio and 0:1 ratio as baseline. (C) Assignment of residues with high chemical shift perturbation upon Ca^2+^ addition and likely direct Ca^2+^ binding residue S491 to existing p6^Gag^ structure (PDB: 2C55). AlphaFold 3 predicts Ca^2+^ to be positioned near p6^Gag^ S491 for possible direct interaction. Average pLDDT are >70. (D) Comparison of late-domain containing sequences (underlined) from ESCRT-dependent viruses, highlighting Ca^2+^ binding (red) and phosphorylation-prone (orange) residues (modified from ^15^). (E) Cartoon illustrating interactions between Ca^2+^ cation and negatively charged/polar amino-acids (in red) or phospho-residues (in orange). (F) Phospho-proteomics discovery of p6^Gag^ and p6^Pol^ phosphorylation sites. Large quantities of cell-free viruses from NL4.3 and BaL strains were processed, and phospho-peptides were positively selected for mass spectrometry (MS) analysis. Summary of phosphorylation sites in p6^Gag^ and p6^Pol^ confirmed/discovered in this study and their prevalence in each strain amongst p6^Gag^ and p6^Pol^ phospho-sites. MS spectra display direct evidence of phosphorylation in p6^Gag^ T456, S488 and S491 and p6^Pol^ S449, T453 and S457.

### Phosphorylation of Pr55^Gag^ and Pr68^GagPR^ modulates Ca^2+^ binding and homo-dimerization of HIV precursor proteins

Building upon earlier works showing virion packaging of mitogen-activated protein kinase 1 (MAPK1/ERK-2) ^21,23^ and *in vitro* phosphorylation of HIV Gag domains with recombinant active extracellular signal-regulated kinase 2 (ERK-2) ^17^, recombinant-Pr55^Gag^ ^24,25^ and - Pr68^GagPR^ (a truncated version of Pr160^GagPol^ used as surrogate) ^8^ were produced and phosphorylated *in vitro* using recombinant ERK-2 for binding studies (***Fig.2***, ***Fig.S3***). Phospho-proteomics confirmed *in vitro* phosphorylation of Pr55^Gag^ on p6^Gag^-S451, -T456, - S488 (***Fig.S3***). Surface plasmon resonance (SPR) analyses showed that phosphorylated Pr55^Gag^ (phospho-Pr55^Gag^) exhibited a 23% decrease in Ca^2+^ binding with dissociation constant (K_D_) increased from 65 ± 5 μM to 84 ± 3 μM (p<0.05) (***Fig.2A-B***). In contrast, phosphorylation of Pr68^GagPR^ (phospho-Pr68^GagPR^) enhanced Ca^2+^ binding affinity by 29% (with K_D_ decreased from 40 ± 5 μM to 31 ± 5 μM, p<0.01) (***Fig.2A-B***). These data imply that phosphorylation of Pr55^Gag^ could have altered the local conformation of p6^Gag^ to reduce its capacity to interact with Ca^2+^, while phosphorylation of Pr68^GagPR^ strengthened its Ca^2+^ binding capacity via local structural modifications.

We next evaluated the impact of phosphorylation on the homo-dimerization of either Pr55^Gag^ (***Fig.2C-E***) or Pr68^GagPR^ (***Fig.2F-H***). The enhanced Ca^2+^-dependent SPR interaction between immobilised Pr55^Gag^ and flowed phospho-Pr55^Gag^ (from K_D_ = 22 ± 6 μM to 8 ± 2 μM, p<0.05, ***Fig.2D***) was not detected in the reverse interaction, where phospho-Pr55^Gag^ was immobilised and Pr55^Gag^ was flowed (***Fig.2C-E***). In contrast, dimerization of phospho-Pr68^GagPR^ in the presence of Ca^2+^ (K_D_ = 0.6 ± 0.3 μM, p<0.0001, ***Fig.2D***) consistently exhibited stronger binding affinity over: (i) Ca^2+^ free dimerization of phospho-Pr68^GagPR^ (K_D_ = 11 ± 2 μM, ***Fig.2D***); (ii) dimerization of phospho-Pr68^GagPR^ / Pr68^GagPR^ with or without Ca^2+^ (***Fig.2F-H***); and (iii) dimerization of non-phosphorylated Pr68^GagPR^ / Pr68^GagPR^ with or without Ca^2+^ (***Fig.2F-H***).

Our *in vitro* analysis of Pr55^Gag^ and Pr68^GagPR^ hetero-dimerization upon phosphorylation have shown that phospho-Pr68^GagPR^ mostly did not negatively impact on *in vitro* SPR interaction with Pr55^Gag^ both in the presence or absence of Ca^2+^ (***Fig.S4***). In comparison, there was a general reduction of binding of phospho-Pr55^Gag^ / Pr68^GagPR^ pair over the binding of Pr55^Gag^ / Pr68^GagPR^ pair (***Fig.S4***). Our *in vitro* SPR analyses suggested: (i) phosphorylation of HIV precursor protein could alter its ability to bind to Ca^2+^ and its interaction with each other; and (ii) Ca^2+^-PO_4_^3-^ bridge could have a role in stabilising HIV precursor protein complexes during particle formation. However, specific cases of Ca^2+^-PO_4_^3-^ functional interaction in HIV replication are needed to demonstrate their roles in biology.

**Figure 2:**
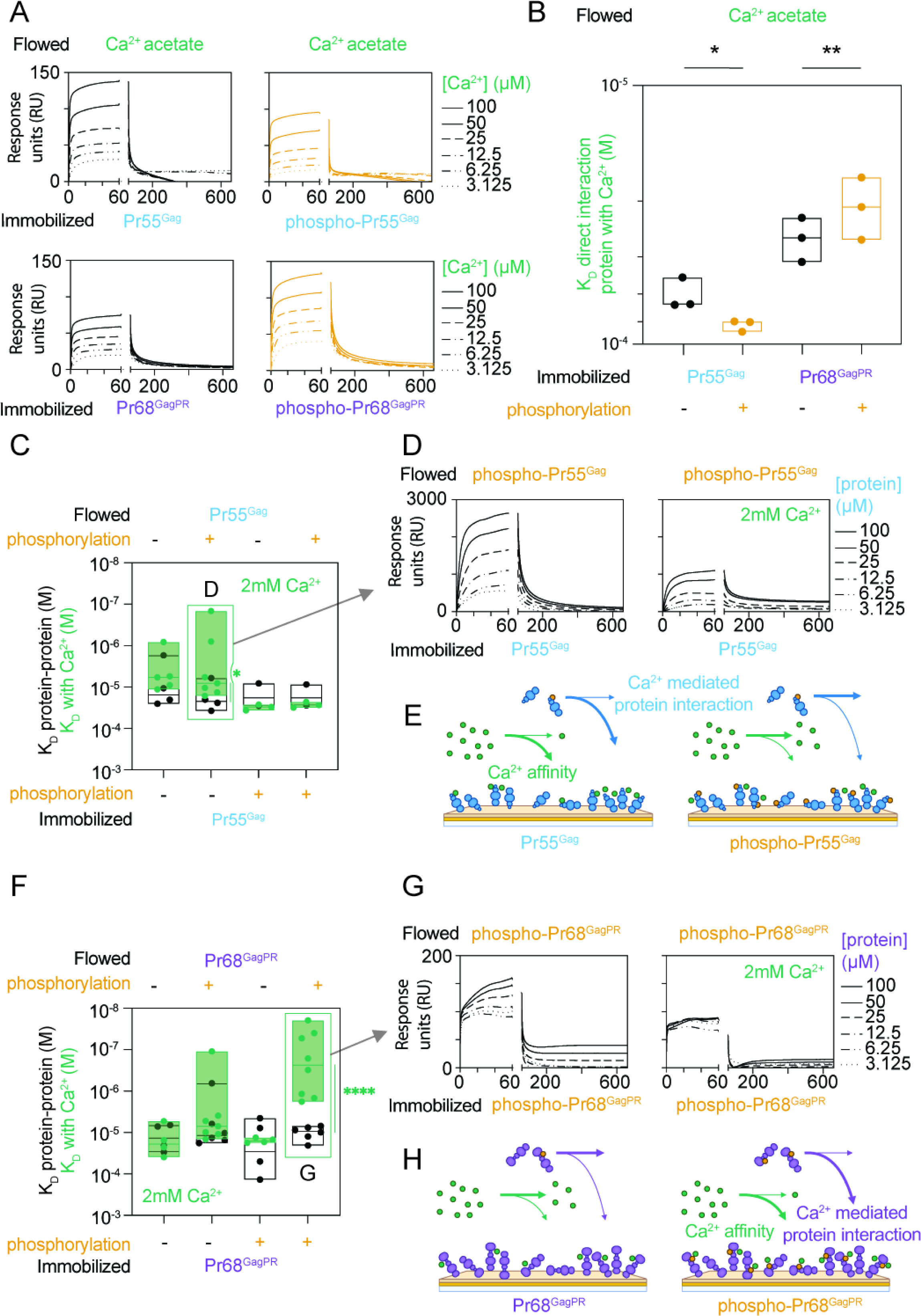
Phosphorylation modulates Ca^2+^-mediated dimerization of HIV proteins. (A) SPR interaction curves of Ca^2+^ flowed at different concentrations (2-fold dilutions from 100 μM to 3.125 μM) with immobilized (*in-vitro* phosphorylated)-Pr55^Gag^/Pr68^GagPR^. (B) Phosphorylation influence on K_D_ of protein-Ca^2+^ interactions. (C) Ca^2+^ enhancement (green vs black) of immobilized (phospho)-Pr55^Gag^ protein-protein interactions with (phsopho)-Pr55^Gag^. (D) SPR interaction curves corresponding to phospho-Pr55^Gag^ flowed against Pr55^Gag^, with or without Ca^2+^, in figure 2C. (E) Model describing Pr55^Gag^ phosphorylation inhibiting Ca^2+^ mediated Pr55^Gag^ protein-protein interactions. (F) Ca^2+^ enhancement (green vs black) of immobilized (phospho)-Pr68^GagPR^ protein-protein interactions with (phsopho)-Pr68^GagPR^. (G) SPR interaction curves corresponding to phospho-Pr68^GagPR^ flowed against phospho-Pr68^GagPR^, with or without Ca^2+^, in figure 2F. (H) Model describing Pr68^GagPR^ phosphorylation promoting Ca^2+^ mediated Pr68^GagPR^ protein-protein interactions. Statistical analysis: paired t-test.

### Evidence of Ca^2+^-PO_4_^3-^ bridge near the C-terminus of p6^Gag^ contributing to HIV function

Our integrated (NMR-, bioinformatic-, and phospho-proteomic-) analyses have implied that phosphorylation capable amino acids of p6^Gag^ (S488, S491, S498, and S499, ***Fig.1***) could be involved in HIV-Ca^2+^ mediated interactions. A combination of biophysical-, molecular virology- and ultra-structural-studies were then employed to identify specific cases of Ca^2+^-PO_4_^3-^ bridge during viral replication and their functional impacts (***Figs.3-5***). In the absence of site-specific phosphorylated recombinant Pr55^Gag^ and Pr68^GagPR^ (surrogate of Pr160^GagPol^), synthetic peptides of p6^Gag^ and p6^Pol^ were used as proxy for SPR analyses (***Figs.3-5***). Despite the controversy about a role of p6^Gag^ phosphorylation in HIV ^17,18,20^, the field agrees that replacement of S/T residues with D/N amino acids (that are capable of Ca^2+^ binding) restores HIV function ^18,20^.

As p6^Gag^ S488 is involved in virion maturation, morphogenesis, and infectivity ^18,26,27^, we first focused on the potential of a p6^Gag^ S488 Ca^2+^-PO_4_^3-^ bridge near the C-terminus of p6^Gag^ to facilitate Pr55^Gag^-Pr55^Gag^ or Pr55^Gag^-Pr160^GagPol^ complex formation (***Fig.3A***). SPR analyses showed phosphorylation of p6^Gag^ S488 (p6^Gag^ pS488) abolished its capacity to: (i) homo-dimerize with p6^Gag^ (***Fig.3B***); and (ii) dimerize with p6^Pol^ in the absence of Ca^2+^ (***Fig.3B***). These data imply that the phosphorylation switch of p6^Gag^ ^pS488^ modulates local p6^Gag^ structures to impact on both Pr55^Gag^-Pr55^Gag^- and Ca^2+^ free Pr55^Gag^-Pr160^GagPol^-interaction (***Fig.3B***). Remarkably, the capacity of p6^Gag^ ^pS488^ to interact with p6^Pol^ was restored in the presence of Ca^2+^ (***Fig.3B***), illustrating a direct interaction between Ca^2+^ and p6^Gag^ pS488 forming a bridge between Pr55^Gag^ and Pr160^GagPol^ during HIV assembly. Importantly, these data have resolved a long-held enigma ^17,18,20^ and demonstrate a functional role of phosphorylation with p6^Gag^ S488 in HIV.

SPR analyses of p6^Gag^ (phospho-mimics, incorporating S488D/S491D/S498D/S499D) were carried out to estimate the relative contributions of other C-terminus p6^Gag^ phosphorylation sites (S491, S498, S499, ***Fig.1E***) to HIV biology in comparison with p6^Gag^ S488. As both p6^Gag^ (S488D/S491D/S498D/S499D) (***Fig.3C***) and p6^Gag^ (pS488) (***Fig.3B***) exhibited similar SPR binding properties, these data implied p6^Gag^-S488 could act as the primary C-terminus p6^Gag^ Ca^2+^-PO_4_^3-^ bridge to stabilize Pr55^Gag^-Pr160^GagPol^ complex, while p6^Gag^ (pS491, pS498, and pS499) could contribute to this p6^Gag^ Ca^2+^-PO_4_^3-^ bridge function in the absence of (pS488) (***Fig.S2***).

Since the AlphaFold 3-predicted Ca^2+^ binding site near p6^Gag^-S491 has been confirmed by NMR-analysis (***Fig. 1B-C***), a combination of SPR and site-directed mutagenesis analyses were done to examine the combined effects of p6^Gag^-S488 and -S491 in HIV biology. SPR analyses highlighted the establishment of an S488/S491 mediated bridge only with p6^Pol^, in the presence of Ca^2+^ (***Fig.3D***). Either complete inactivation of phosphorylation (p6^Gag^ S488A/S491A) or phosphomimic mutations to simulate constitutive phosphorylation (p6^Gag^ S488D/S491D) resulted in reduction of HIV infection in primary cells (***Fig.3E***). The functional defects of HIV (S488A/S491A) and HIV (S488D/S491D) (***Fig.3E***) were neither related to the proteolytic processing of HIV proteins during maturation (***Fig.3F***) nor associated with the virion packaging of Pr160^GagPol^ precursor during HIV assembly (***Fig.3G***). Consistent with our virion Phospho-proteomics analysis (***Fig.1F***), these data illustrated that HIV only requires a fraction of p6^Gag^ S488 (and/or C-terminus p6^Gag^, ***Fig.S2***) to be phosphorylated (or transiently phosphorylated) for the formation of Ca^2+^-PO_4_^3-^ bridge, while excessive (or complete) phosphorylation of S488 is detrimental to HIV function that is consistent with transient modulation / switching mechanism in biology.

**Figure 3:**
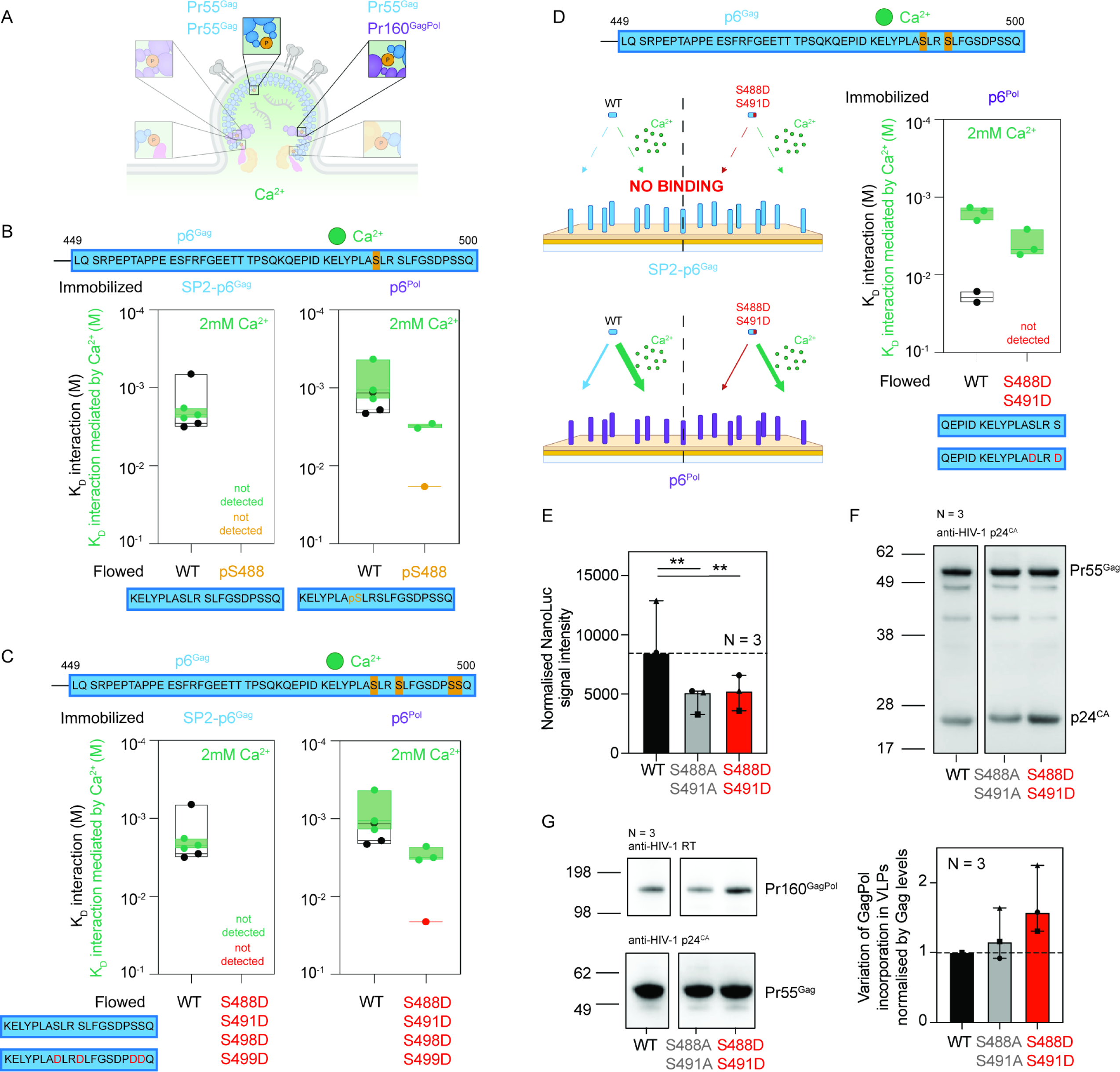
p6^Gag^-S488 and -S491 form Ca^2+^-PO_4_^3-^ bridge to support HIV assembly. (A) Focus on Ca^2+^ mediated (phospho)-proteins interactions including Pr55^Gag^-Pr55^Gag^ and Pr55^Gag^-Pr160^GagPol^ during HIV assembly. (B, C, D) SPR measurements of fragment interaction K_D_ with immobilised SP2-p6^Gag^/p6^Pol^ in presence or absence of Ca^2+^. (B) WT fragment is compared to pS488 phospho-peptide. (C) WT fragment is compared to S488D/S491D/S498D/S499D phospho-mimic. (D) WT fragment is compared to S488D/S491D phospho-mimic. Interactions with SP2-p6^Gag^ are not detected. (E) Relative infectivity of HIV S488A S491A and HIV S488D S491D particles compared to WT. WT control conditions are identical to those used for comparison with (non)-phospho-mimicking conditions in Figure 5D. Statistical analysis: repeated measures one-way ANOVA. (F) Proteolytic processing patterns of mutant particles mimicking phosphorylation states compared to WT. (G) Phosphorylation mimics effect on Pr160^GagPol^ packaging levels in immature particles.

### Ca^2+^-PO_4_^3-^ bridge between p6^Pol^ domains could promote Pr160^GagPol^ dimerization for virion maturation

The phosphorylation patterns (***Fig.S2)*** and functional impacts of p6^Gag^ S488, S491, S498, and S499 (***Fig.3*)** would suggest redundancy exists within a specific group of putative regulatory amino acids. Our phospho-proteomic study has identified five novel phosphorylation sites in p6^Pol^ (p6^Pol^ S449, S450, T453, S457, and T459) (***Fig.1F, S2***) which could result in 32 different phosphorylation variants containing from zero to five phosphorylated S/T within a single p6^Pol^. In the absence of functional data about any p6^Pol^ amino acid phosphorylation, we analysed these five identified p6^Pol^ phosphorylation amino acids (***Fig.S2***) as a group. AlphaFold 3 ^14^ prediction illustrated Ca^2+^-mediated p6^Pol^ complex could be established between phosphorylated p6^Pol^ dimer (pS449/pS450/pT453/pS457/pT459) and phospho-mimic p6^Pol^ dimer (S449D/S450D/T453D/S457D/T459D) at reasonable levels of confidence with pLDDT > 70 (***Fig.4A, S5***). By focusing on the functional contribution of Ca^2+^-PO_4_^3-^ bridge to Pr160^GagPol^ dimerization (***Fig.4B***), SPR analyses showed phospho-mimic p6^Pol^ peptides abolished capacity of p6^Pol^ to dimerize in the absence of Ca^2+^ (from K_D_ = 2.4 ± 0.3 mM to undetectable, ***Fig.4C***) but enhanced the binding affinity 7 times compared to WT in the presence of Ca^2+^ (from K_D_ = 2.8 ± 0.2 mM to 0.4 ± 0.2 mM, p<0.01, ***Fig.4C***). SPR data were consistent with AlphaFold 3 predictions that phosphorylation / phospho-mimic could lead to local structural changes to prevent p6^Pol^ dimerization in the absence of Ca^2+^, but Ca^2+^-mediated interaction restored complex formation (***Fig.4B-C***). The ability of phospho-mimic p6^Pol^ peptides (S449D/S450D/T453D/S457D/T459D) to dimerize only in the presence of Ca^2+^ demonstrated a direct-interaction between Ca^2+^ and bidentate ligands from either phosphomimic amino acids (D) or phosphorylated S/T to stabilise protein complex via Ca^2+^-PO_4_^3-^ bridge (***Fig.4C***).

Analyses of sites-directed HIV mutants with: (i) phosphorylation inactivation in p6^Pol^ (HIV p6^Pol^ non-phospho-mimic S449A/S450A/T453A/S457A/T459A); and (ii) phosphomimic mutations to simulate constitutive phosphorylation in p6^Pol^ (HIV p6^Pol^ phospho-mimic S449D/S450D/T453D/S457D/T459D), have shown that neither virion packaging of Pr160^GagPol^ (***Fig.4D***) nor proteolytic processing of virion proteins (***Fig.4E***) was impacted by the phosphorylation status of p6^Pol^. However, an enhancement of viral infectivity was detected with p6^Pol^ phospho-mimic HIV (***Fig.4F***). Transmission electron microscopy (TEM) analyses showed newly released HIV p6^Pol^ phospho-mimic particles often associated with greater number of virions with electron dense cores (***Fig.4G***), possibly indicating an alternative stage of virion maturation upon release. Using this hyper-phosphorylated p6^Pol^ mimic mutant, our work suggested Ca^2+^-PO_4_^3-^ bridge in p6^Pol^ phosphorylation sites could play a role in the dimerization of HIV Pr160^GagPol^ and facilitate the virion maturation process during particle assembly and release.

**Figure 4:**
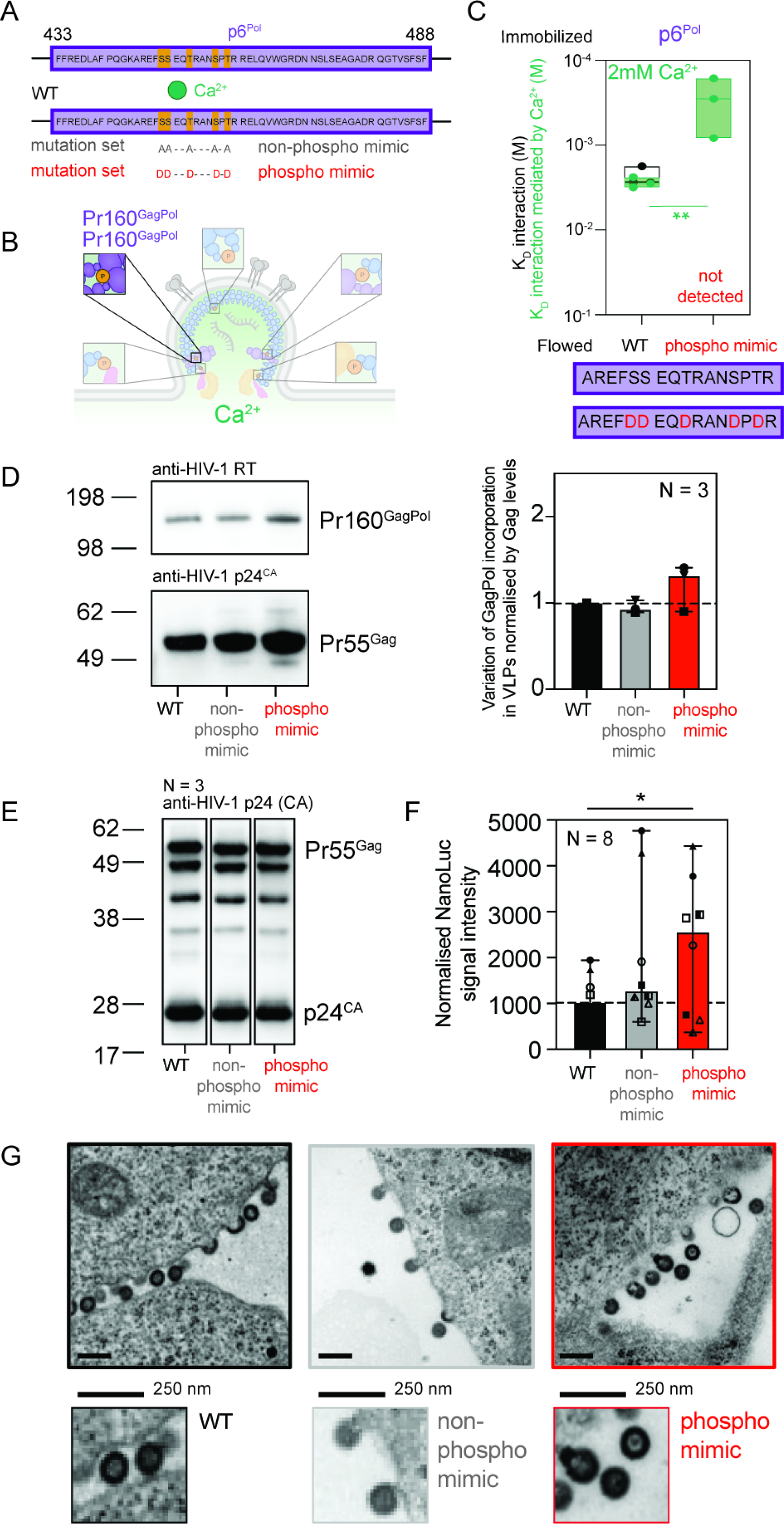
Ca^2+^-PO_4_^3-^ bridge potentiates HIV maturation by facilitating p6^Pol^ dimerization. (A) Engineered mutations within p6^Pol^ to mimic non-phosphorylated sites (non-phosphomimic mutations S/T to A) or phosphorylated sites (phosphomimic mutations S/T to D). (B) Studied Ca^2+^ mediated (phospho)-proteins interactions during HIV assembly. (C) SPR analysis of WT / phospho-mimic peptides interactions with immobilised p6^Pol^ in presence or absence of Ca^2+^. Statistical analysis: unpaired t-test (D) Phosphorylation mimics effect on Pr160^GagPol^ packaging levels in immature particles. (E) Proteolytic processing patterns of mutant particles mimicking both p6^Pol^ phosphorylation states compared to WT pattern. (F) Efficiency of p6^Pol^ phospho-mimic particle genome delivery efficiency compared to WT. Statistical analysis: repeated measures one-way ANOVA. (G) TEM imaging of early budding particles (WT and phospho-mimics) and their electron density distribution. Captions: WT/phospho-mimic particles and their electron density distribution enlarged 2x.

### Virion packaging of enzymatic proteins and cellular factors modulation via a PTAP p6^Gag^ T456 Ca^2+^-PO_4_^3-^ bridge

All ESCRT binding motifs have a putative phosphorylation site ^15^ (***Fig.1D***). Our work has confirmed the phosphorylation status of HIV ESCRT binding motifs (***Fig.1E, S2***). AlphaFold 3 was used to simulate the interaction between p6^Gag^ PTAP containing fragment and TSG101 ubiquitin E2 variant (UEV) domain in the presence of Ca^2+^ (***Fig.5A, S6***). Upon phosphorylation of p6^Gag^ T456 (p6^Gag^ pT456), p6^Gag^ pT456 was predicted with high confidence to be oriented away from TSG101 UEV for Ca^2+^ interaction (***Fig.5A, S6***), implying a potential regulatory role of p6^Gag^ T456 to complex with other proteins during assembly (***Fig.5B***). Our SPR analyses showed phospho-mimic at p6^Gag^ S451 T456 (p6^Gag^ S451D T456D) eliminated p6^Gag^-p6^Gag^ interaction and Ca^2+^ free p6^Gag^-p6^Pol^ interaction (***Fig.5C***). However, p6^Gag^ S451D T456D retained the capacity to support complex formation of p6^Gag^-p6^Pol^ in the presence of Ca^2+^ (***Fig.5C***). These data are consistent with a role of p6^Gag^ T456 phosphorylation in forming a direct interaction with Ca^2+^ (i.e. Ca^2+^-PO_4_^3-^ bridge) to stabilise Pr55^Gag^-Pr160^GgaPol^ complex for virion packaging of the HIV enzymatic precursor proteins (***Fig.5A***).

To delineate the precise functional contribution of p6^Gag^ pT456 during HIV assembly, we analysed the impacts of site-directed HIV mutants via: (i) phosphorylation inactivation in p6^Gag^ T456 (p6^Gag^ T456A); and (ii) phosphomimic mutations to simulate constitutively phosphorylated p6^Gag^ T456 (p6^Gag^ T456D). Both HIV T456A and HIV T456D exhibited a 50% reduction of infectivity in comparison with wild type (***Fig.5D***), indicating partial phosphorylation of p6^Gag^ T456 is important for HIV to function. Western blot analyses of HIV particles showed greater abundance of p25^CA-SP1^ in HIV T456A than HIV T456D, which in turn was greater than wildtype control (***Fig.5E***). These data demonstrated defects in maturation / proteolytic processing for these p6^Gag^ T456 phosphorylation dysregulated mutants.

Comparison of virion packaging of Pr160^GagPol^ revealed a 50% suppression of viral enzyme incorporation in HIV T456A, whilst no significant alteration of Pr160^GagPol^ packaging was detected with HIV T456D (***Fig.5F***). The observed defect of enzyme packaging of HIV T456A (***Fig.5F***) confirmed p6^Gag^ T456 Ca^2+^-PO_4_^3-^ bridge in stabilising Pr55^Gag^-Pr160^GagPol^ complex during HIV assembly. Analyses of ESCRT protein packaging in virions showed that both HIV T456A and HIV T456D exhibited defects in TSG101 packaging, with greater repression seen with HIV T456A (***Fig.5G***). The virion-associated ALIX protein in both HIV T456A and HIV T456D being equivalent to wildtype (***Fig.5G***) highlighted the p6^Gag^ pT456 induced structural impacts were locally confined, which was consistent with AlphaFold 3 prediction (***Fig.5A***). TEM analyses confirmed both HIV T456A and HIV T456D displayed classical HIV-1 late domain PTAP mutation phenotypes (***Fig.5H***) ^28^. Together, these data illustrate that p6^Gag^ pT456 Ca^2+^-PO_4_^3-^ bridge functions as a dual modulator of Pr160^GagPol^ virion packaging and of direct interactions with TSG101 for particle release.

**Figure 5:**
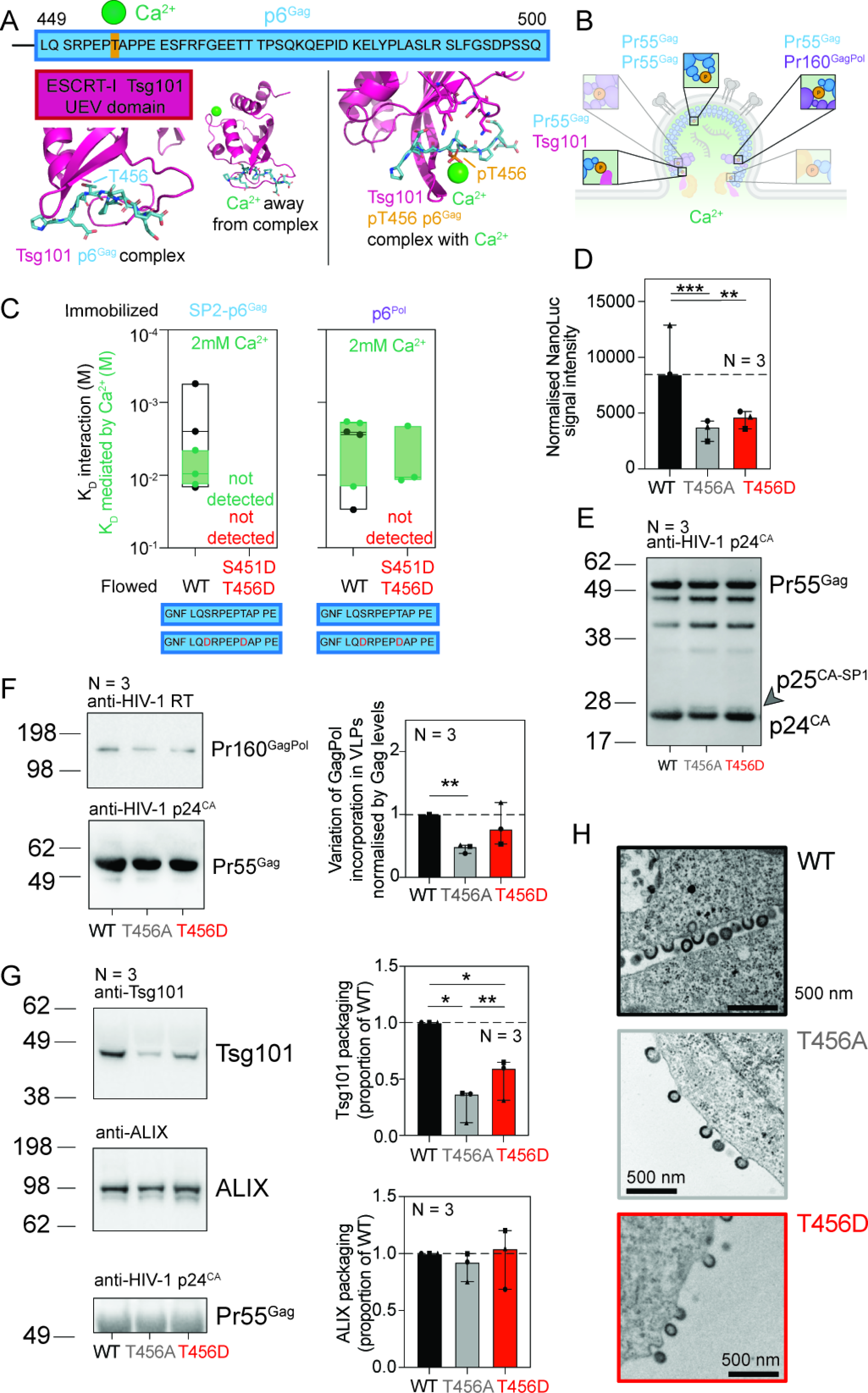
A PTAP Ca^2+^-PO_4_^3-^ bridge regulates particle assembly and release. (A) Structure prediction of Tsg101 UEV domain complex with PTAP containing p6^Gag^ fragment in presence of Ca^2+^ (based on PDB: 1M4P) ^29,30^ has predicted structure of ipTM = 0.74 and pTM =0.91. AlphaFold 3 also predicted phosphorylation of T456 in p6^Gag^ would push T456 side chain away from UEV and to interact with Ca^2+^ ion, with ipTM = 0.79 and pTM = 0.90. p6^Gag^ with or without Ca^2+^ and T456 phosphorylation. (B) Protein-protein interaction studies of p6^Gag^ T456 during complex formation. (C) SPR analysis of WT / phospho-mimic peptide interactions with immobilised SP2-p6^Gag^/p6^Pol^ in the presence and absence of Ca^2+^. (D) Relative infectivity comparison between HIV p6^Gag^ T456 mutants (HIV T456A and HIV T456D) against WT control. Statistical analysis: repeated measures one-way ANOVA. (E) Pr55^Gag^ proteolytic processing pattern of WT and phospho-mimic particles. Incomplete processing results in a double band at 25/24 kDa, indicated by an arrow. (F) Comparison of Pr160^GagPol^ packaging levels between WT and phospho-mimic immature particles. Statistical analysis: paired t-test. (G) Cellular Tsg101 and ALIX packaging levels in phospho-mimic particles compared to WT. WT control conditions are identical to those used for comparison with (non)-phospho-mimicking conditions in Figure 3F. Statistical analysis: paired t-test. (H) TEM imaging of early budding particles (WT and phospho-mimics) illustrating their Tsg101-mediated budding capacity.

## DISCUSSION

Coordination of proteins complex formation is of major importance in biology. A significant portion of Ca^2+^ signalling or phosphorylation switch is to regulate the interactions between biological complexes. Recognising the shared properties between Ca^2+^ binding amino acids (D, E, N, & Q) and phosphorylated amino acids (pS, pT, & pY), we have now identified phosphorylation switching has a second layer of control via Ca^2+^-mediated complex formation, illustrating how two distinct branches of cell biology mechanisms work cooperatively to support biological functions. Specifically, our data demonstrate that Ca^2+^-stabilised protein complexes could be achieved via the formation of Ca^2+^-PO_4_^3-^ bridge(s) across molecules, representing a previously unknown mechanism in Ca^2+^ signalling and phosphorylation switch biology ^2,4^.

A critical aspect of any regulatory system is quality control to maintain system efficiency. We have previously shown that HIV relies on interactions with Ca^2+^ derived from intracellular Ca^2+^ release units (CRUs) to achieve directional uropod targeting of HIV assembly complex and virological synapse formation ^8^, and disrupting these Ca^2+^-based interactions can lead to ubiquitination and degradation of HIV proteins. Building upon accumulated knowledge in Ca^2+^ spark biology ^3^, we proposed a model where numerous CRUs across the cell function as a general cellular digital traffic control system facilitating intracellular trafficking ^8^. Consequently, interfering with Ca^2+^-mediated complex formation would lead to premature protein degradation via ubiquitination ^8^. By extension, should Ca^2+^-mediated (and/or Ca^2+^-PO_4_^3-^ bridge stabilised) complex formation ^8^ be a general mechanism, corroborative evidence must exist in published literature supporting this hypothesis.

*First,* the concept of a relationship between Ca^2+^ and phosphorylation is not new. An interplay between Ca^2+^ and phosphorylation was described in the original phosphorylation switch paper ^31^ ahead of the discovery of Ca^2+^ sparks in heart muscle cells ^32^. *Second,* evolutionary driven convergent enrichment of: (i) Ca^2+^ binding amino acids; and (ii) putative phosphorylation sites, flanking ESCRT binding motifs across diverse viral genomes illustrates the functional significance of this relationship. *Third,* a verified phosphorylation sites prediction model (DISPHOS) ^33^ has identified that Ca^2+^-binding amino acids are statistically enriched near phosphorylation sites ^33^. Interestingly, DISPHOS was established, in part, based on ‘protein phosphorylation sites’ ^33^ consisting of both ‘disordered structures’ and ‘proline residues’, which are also well-known features found in ‘regions surrounding viral ESCRT binding motifs’ ^15,28^. *Fourth,* our observation that ubiquitination acts as a quality control checkpoint inducing premature degradation of dysfunctional Ca^2+^-facilitated complexes ^8^ is in practice similar to the described PEST sequence mediating protein degradation ^34^. Importantly, the reported PEST sequences are enriched with both putative phosphorylation sites and Ca^2+^-binding amino acids ^34^, analogous with sequences surrounding the ESCRT binding motifs in viral sequences. *Fifth,* if the interplay between Ca^2+^ and phosphorylation switch is a general principle, other cellular examples must exist. In the context of exocytosis biology, we noticed that sequences surrounding key phosphorylation switch determinants of exocytosis-related trafficking proteins, such as Exo70 ^35^, and Synapsins (I, II, and III) ^36^ are enriched in Ca^2+^-binding amino acids ^35,36^. Reciprocally, the Ca^2+^ binding amino acids on exocytosis related protein, Synaptotagmin I ^37^, are surrounded by multiple putative phosphorylation sites ^37^.

As viruses navigate the host cell system in a myriad of ways to support their own survival, viruses offer an exceptional vantage point to discover unknown cellular processes. The ‘simplicity’ of virus genomes dictates viruses to exploit existing cellular processes to overcome ‘barriers’ against viral propagation. Using Ca^2+^-dependent polarised targeting of HIV as a model system, our work identified an unexpected layer of regulation that bring Ca^2+^ signalling and phosphorylation switch under one roof. Understanding the contribution of Ca^2+^-PO_4_^3-^bridge in the regulation of other phosphorylation switch biology has the potential to identify unconventional approaches in translational science.

## Supporting information

Supplementary Data for Bremaud et al

## ACKNOWLEDGEMENTS

General

We express our gratitude to both Julian W. Bess and Jeffery D. Lifson at Leidos Biomedical Research Inc., Frederick National Laboratory, for Cancer Research, USA, for generously providing us with large quantity of purified HIV for virion phospho-proteomic analyses. These precious reagents are the foundation of both our discovery and data presented in this manuscript. The phosphoproteomic analysis of HIV particles corresponds toa fee-for-service arrangement at Children’s Hospital of Philadelphia Research Institute, Philadelphia, Pennsylvania, USA. We thank Markus Thali, Benjamin Chen, and Owen Pornillos for their insightful suggestions during data discussions.

## Author Contributions

EB, BLS, and JM designed the study. EB, BPM, AB, VM, TF, AE-D, LH-T, and BLS performed the experiments. EB and JM wrote the manuscript. All authors contributed to the discussions on the experiments, the collected data, and the finalisation of manuscript.

## Data and materials availability

All data needed to evaluate the conclusions in the paper are present in the paper and / or the Supplementary Materials. Additional data related to this may be requested from the authors.

## MATERIALS AND METHODS

### Structure predictions

AlphaFold3 (https://alphafoldserver.com) was used for modelling. PyMOL was used to analyse the models and generate annotated schematics.

### Peptides

Isotope labelled peptides used in NMR assays were purchased from Mimotopes Pty Ltd (Australia). Peptides used in SPR assays were purchased from GenScript Biotech (USA).

### Plasmids

All plasmids were engineered by standard molecular biology techniques, through plasmid restriction and gBlock (Integrated DNA Technologies) recombination using NEBuilder® HiFi DNA Assembly (New England Biolabs, E5520S). Generated plasmid sequences were validated by capillary electrophoresis sequencing through Macrogen CO., Ltd (S. Korea).

In brief, site-directed mutagenesis studies were achieved by decoupling overlapping Gag and GagPol sequences, and non-frameshifting Gag (or mutant) and frameshifting GagPol (or mutant) constructs were co-transfected in a 20 to 1 ratio to mimic observed frameshifting frequency. Frameshifting was blocked in Gag construct through replacing the slippery sequence nucleotides TTTTTTA in positions 1296-1333 with CTTCTTC. On the contrary, frameshifting was forced in GagPol constructs through replacing this same slippery sequence with CTTCTTCA. In applicable conditions, HIV protease was inactivated by site-directed D25N mutation in PR.

### Recombinant protein expression

Recombinant protein constructs were expressed in *Escherichia coli* BL21AI strain, cultured at 37°C in Terrific Broth supplemented with 17 mM monopotassium phosphate (KH_2_PO_4_) and 72 mM dipotassium phosphate (K_2_HPO_4_), 50 μM of zinc sulphate (ZnSO_4_) and 30 μg/mL kanamycin. When the cultures’ OD_600_ ranged between 0.6 and 0.8, protein production was induced by adding 1 mM Isopropyl ß-D-1-thiogalactopyranoside (IPTG) and 0.1 mM arabinose. Cells expressing Pr55^Gag^ TEV-His or Pr68^GagPR^ TEV-His were cultured for 20 hours at 180 rpm, 18°C, whereas cells expressing p15 NC-SP2-p6^Gag^ TEV-His or p15 NC-p6^Pol^ TEV-His were maintained for 4 hours at 180 rpm, 37°C. Cultures were harvested by centrifugation at 5000 g for 10 min at 4°C, and pellets were frozen at -80°C.

### Recombinant protein purification

Bacterial pellets from 2 L of cultures were lysed in 80 mL of lysis buffer (50 mM TRIS-base pH 8.0, 1.0 M NaCl, 5 mM MgCl_2_, 10 mM imidazole, 10 % (v/v) glycerol, 1.0 % (v/v) Tween-20, 1 mM phenylmethylsulfonyl fluoride (PMSF), 1 mM benzamidine-HCl, 5 mU benzonase, 0.5 mg/mL lysozyme, 0.5 mM tris(2-carboxyethyl)phosphine (TCEP)). The cell lysate was homogenised using a cell disruptor at 18,000 psi (Constant Systems). Lysates were then clarified by centrifugation at 15,000 *g*, 4°C for 30 min. His-tag proteins were purified by capture chromatography using AKTA Start with HisTRAP FF 5 mL column (GE Lifesciences). The C-terminal poly-his tag was then cleaved by using recombinant TEV protease in SnakeSkin Dialysis Tubing (ThermoFisher, 68035) with overnight dialysis at 4°C in SEC Buffer (50 mM TRIS-base pH=8.0, 1.0 M NaCl, 1 mM TCEP). Cleaved poly-His was separated from proteins of interest by capture chromatography with HisTRAP FF and proteins were subsequently concentrated using appropriate Ultracel® regenerated cellulose Amicon Ultra 15 centrifugal filters. Proteins were separated in size-exclusion chromatography (SEC) Buffer by polishing size exclusion chromatography using GE Lifesciences columns HiLoad 26/600 Superdex 200 pg (alternatively 75 pg Increase for 15 kDa proteins). Relevant fractions were pooled, concentrated using Amicon Ultra 15 centrifugal filters and frozen at -80°C.

### *In-vitro* phosphorylation assays

Recombinant active, GST tagged, human ERK-2 (Sigma Aldrich, E1283-10UG) was used for *in-vitro* phosphorylation of purified HIV-1 Gag derivatives Pr68^GagPR^ or Pr55^Gag^. The enzyme was used at a concentration 5-20 times greater than required to match the batch specific activity. Recombinant Pr68^GagPR^ / Pr55^Gag^ (400 μg) were introduced in 10 mM Tris-HCl (pH 7.5), 5 mM MgCl_2_, 0.5 mM dithiothreitol (DTT) with the addition of 1.25 – 5 mol ATP and 1.11 – 4.44 μg ERK-2. The reaction was maintained at 30°C for 60min, subsequently, elution through GSTrap^TM^ FF column (17513002, Cytiva) stopped the reaction. Flowthrough containing phosphorylated recombinant proteins was collected, A_280_ was measured using NanoDrop 2000/2000c and used to calculate protein concentration, and stocks were aliquoted and frozen at -80°C.

### Surface plasmon resonance of (phospho)-proteins

The proteins (phospho)-Pr55^Gag^ and (phospho)-Pr68^GagPR^ were immobilized onto Series S CM5 chip activated surfaces. Briefly, using GE Healthcare BIAcore S200/T200 instruments, the proteins were captured using amine coupling, in which the carboxylmethyl dextran matrix of the sensor chip was activated by a 720 sec injection of a mixture of 0.2 M 1-ethyl-3-[(3-dimethylamino)propyl]-carbodiimide (EDC) and 0.05 M N-hydroxysuccinimide (NHS), which resulted in the conversion of carboxyl groups to NHS esters. The proteins were diluted in 10 mM sodium acetate buffer pH 4.0 at a concentration of 100 µg/mL and immobilized at levels ranging between 9000-12000 RU on flow cells 2-4. The remaining unreacted NHS ester groups were neutralized by an injection of 1 M ethanolamine-HCl (pH 8.0). Flow cell 1 was a negative control, where the flow cell underwent the same treatment as the active flow cells, without protein immobilization. This enabled reference cell subtraction of the responses (2-1, 3-1, 4-1).

Flowed calcium-acetate stock solution was prepared at 10 mM in milliQ water, flowed (phospho)-proteins were diluted in PBS (137 mM NaCl, 2.7 mM KCl, 10 mM Na_2_HPO_4_, 2 mM KH_2_PO_4_), and further diluted in 2-fold dilution series, ranging from 100 µM to 3.125 µM.

Affinity analysis of the binding interaction was performed using multi-cycle model, where ligands were flowed at 30 µL/min rate, with or without calcium proteins-interaction enhancement step using 2 mM calcium-acetate. A series of buffer only controls enabled blank subtraction of the reference subtracted sensograms. A 30 sec injection of 10mM Glycine pH 2.0 at 30 µL/min was used as a regeneration buffer. Data processing and analysis was performed using BIAcore S200/T200 Evaluation Software (Cytiva).

### Surface plasmon resonance of (phospho)-peptides

The biotinylated peptides were immobilized onto a Series S SA (streptavidin) sensorchip. Briefly, using a GE Healthcare BIAcore T200/1S+ instruments, the biotinylated-peptides were captured following sensor chip activation by a 720 sec injection of a 50% isopropanol, 1 M NaCl, 0.1 M NaOH and 1 M NaCl, 0.1 M NaOH. The biotinylated-peptides were diluted to 15 µg/mL and immobilized at levels ranging between 1300-1600 RU. Two flow cells were used as negative controls and underwent the same treatment as the active flow cells, without peptide immobilization. This enabled reference cell subtraction of the responses.

Flowed peptides were reconstructed in HBS-EP (10 mM HEPES, 150 mM NaCl pH 7.4, 0.05% Tween20, 35 mM EDTA) or alternatively HBS-P (no EDTA), and further diluted in 2-fold dilution series, ranging from 200 µM to 12.5 µM.

Affinity analysis of the binding interaction was performed using single-cycle model with or without calcium enhancement step using 2 mM calcium-acetate, where the peptides were flowed at 30 µL/min rate. A series of buffer only controls enabled blank subtraction of the reference subtracted sensograms. A 30 sec injection of 10mM Glycine pH 2.0 at 30 µL/min was used as a regeneration buffer. Data processing and analysis was performed using BIAcore T200/1S+ Evaluation Software (Cytiva).

### Nuclear Magnetic Resonance (NMR) Spectroscopy

^1^H-^15^N heteronuclear single quantum coherence (HSQC) spectra were acquired on a Bruker Avance Neo NMR spectrometer operating at 600 MHz resonance frequency equipped with a quadruple resonance probe. All ^1^H-^15^N HSQC titrations between p6^Gag^ and Ca^2+^ were carried out under identical experimental conditions of 8 scans and 400 complex points in the indirect dimension. The concentration of selectively labelled (Leu, Iso and Phe residues only) ^15^N-p6^Gag^ was kept constant at 80 uM, whereas the Ca^2+^ concentration was gradually increased to 400 µM (1:5). Each individual ^1^H-^15^N spectrum of p6^Gag^ was processed by TopSpin 4.4.0 (Bruker, Massachusetts, USA). and MNova (Mestrelab Research, Spain). Peaks were assigned using the previously solved NMR structure (BMRB entry 15957, PDB ID 2C55) ^13^. Peak assignments were performed in MNova using 2D ^1^H-^1^H TOCSY chemical shifts for p6^Gag^. The chemical shift changes were calculated (equation 1) ^22^, and plotted to characterize the residues undergoing maximum perturbations with increasing Ca^2+^ concentrations (peptide to Ca^2+^ ratios of 0:1, 1:1 and 1:5 were used).

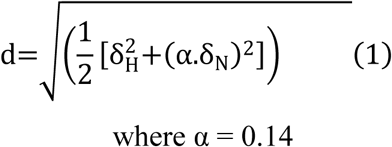

### Cell culture and transfection for lentiviral vector generation

HEK-293T cells were maintained in DMEM supplemented with 10% (v/v) fetal bovine serum (FBS), 100 U/mL penicillin and 100 μg/mL streptomycin. Lentiviral vector production was initiated by co-transfection of Gag and GagPol constructs (+/-mutations) in a 20:1 ratio into HEK-293T. Transfection was carried out with a total of 12 μg plasmid DNA per 7.5x10^6^ cells seeded the day before using PEI-Max (Polysciences Inc.). Supernatants were harvested 48 hours post-transfection, centrifuged and filtered using 0.45 μm cellulose acetate membranes (Sartorius, 16555). Particles were concentrated 700 times by ultracentrifugation using SW 32 Ti swinging-bucket rotor in Beckman Coulter Optima XPN-100 Ultracentrifuge at 26,500 rpm or 86,000 x *g*, for 1 hour at 4°C (ω^2^t = 2.77x10^10^) through 20% (w/v) sucrose cushion. Pellets were resuspended in Dulbecco’s phosphate buffered saline (DPBS) and stored at -80°C. Stocks were quantified using ELISA-p24 kit (XpressBio, XB-1010).

Peripheral Blood Mononuclear Cells (PBMCs) were isolated from fresh concentrated buffy packs (Australian Red Cross), within 24 hours of blood draw ^38,39^. The buffy packs were diluted 1:1.5 with 1X phosphate buffered saline, (pH 7.2, Gibco, 70013-032). Red blood cells were removed by pelleting through a layer of Lymphoprep density gradient medium (Stemcell, 07851) by centrifugation at 800 *g* for 25 min. Cells of interest were then collected at the interface between plasma and LymphoPrep and washed twice by centrifugation at 320 *g* for 10 min at 4°C followed by pellet resuspension in 1X PBS, pH 7.2 to deplete in platelets. Cells were then washed three times by centrifugation at 130 *g* for 10 min at 4°C and ultimately resuspended in complete RPMI containing 10% FBS, 100 U/mL penicillin, 100 µg/mL streptomycin and 2 mM L-glutamine. Cells were incubated at 37°C, 5% CO_2_ for 30 to 45 min in three successive T175 Flasks (Sarstedt, 83.3912.002) to deplete macrophages. PBMCs were seeded at a density of 2x10^6^ cells per mL of complete RPMI supplemented with 50 U/mL human interleukin-2 (hIL-2, Roche, 11147528001) and 10 μg/mL phytohemagglutinin-L (PHA-L, Roche, 11249738001) and activated for 3 to 4 days at 37°C, 5% CO_2_.

### Western blot analysis

Samples were lysed with 1% TritonX-100 (Sigma, T8787) and separated by SDS-PAGE 10% Bis-Tris NuPAGE (Invitrogen, NW00105BOX) and transferred to nitrocellulose membrane (GE Healthcare). Membranes were blocked with 5% (w/v) skim milk in Tris Buffered Saline, 0.05% Tween20 (TBST), washed and probed with either mouse monoclonal anti-p24^CA^ (AG3.0, NIH AIDS Reagent Programme), mouse monoclonal anti-RT (5B2B2, NIH AIDS Reagent Programme), rabbit monoclonal anti-TSG101 (E6V1X, Cell Signalling, 72312) or rabbit monoclonal anti-ALIX (E6P9B, Cell Signalling, 92880). HRP conjugated anti-mouse (Dako, P0161) or HRP conjugated goat anti-rabbit IgG (H+L) (Dako, P0448) were used as secondary antibodies, and blots were imaged by chemiluminescence (SuperSignal TM, Thermo scientific, 34580). Western blots were imaged using BioRad ChemiDoc XRS+.

### Infectivity assays

Lentiviral vectors produced for this assay were pseudotyped with HIV-1 A1 envelope (Env) and packaged a nano-luciferase (NanoLuc) coding sequence. A total of 12 μg plasmids coding for Gag, GagPol (+/-mutations), Env and NanoLuc in a 20:1:2:2 ratio respectively, were used to co-transfect 7.5x10^6^ HEK-293T seeded the day before. Vectors were collected and concentrated as described above.

Activated human PBMCs were seeded in fresh complete RPMI supplemented with hIL-2 and PHA-L at 5x10^6^ cells per condition and then transduced with 500 ng of lentiviral vectors per condition for 2 days at 37°C.

Cells were counted, washed with Hanks’ Balanced Salt Solution (HBSS, Gibco, 14025-092) and lysed in corresponding volumes of lysis buffer (100 mM NaCl, 10 mM Tris-Cl, pH 7.5, 1 mM EDTA, 1% IGEPAL CA-630) to reach 2.5x10^7^ PBMCs/mL lysis buffer.

The efficiency of NanoLuc delivery was measured using Nano-Glo Luciferase Assay System (Promega, N1110) according to manufacturer’s instructions. After 3 minutes, samples were analysed in Costar 96 flat bottom, white plates in a Tecan Infinite M200 Pro Reader (1s integration luminescence).

### Thin section TEM

Transfection of 1x10^7^ HEK-293T cells was achieved in tissue culture treated dishes (Corning, 430167) as described earlier ^39,40^. Cells were fixed after 28h with 2.5% glutaraldehyde in PBS overnight at 4°C. The samples were then washed once with PBS, followed by another wash in PBS using a Pelco Biowave for 40 sec at 250 W at ambient pressure. Post fixation was performed using 1% Osmium tetroxide in a Pelco Biowave’s (2 min power on, 2 min power off, 2 min power on) programmed at 80 W under vacuum and repeating the program without buffer exchange. Post fixed cells were washed once with PBS followed by another wash in PBS using a Pelco Biowave for 40 sec at 250 W at ambient pressure. Cells were dehydrated with a series of increased concentrations of ethanol: 30%, 50%, 70%, 90% and 100% (twice) for 40 sec at 250 W at ambient pressure using a Pelco Biowave. After dehydration, cells were infiltrated with increasing concentration of LX112 resin: first with LX112 resin and ethanol mixture ratio at 1:2 and 2:1 and followed by two infiltrations with 100% resin for 3 min at 250 W under vacuum in a Pelco Biowave. Excess 100% infiltrated resin was removed from the petri dish and 100% LX112 resin filled in BEEM capsule, which were then inverted over the cell dense area of petri dish (3 BEEM capsules prepared for each sample).

Samples were kept in 60°C oven up to 48 hours for resin polymerisation. The BEEM capsules were then snapped off from the petri dish, to achieve smooth surface on resin block. A resin block with a large, smooth surface was selected from each sample for easy alignment of the resin block face to the diamond knife in the later sectioning stage. For sectioning, a Leica ultramicrotome (Ultracut UC71) was used.

The resin block was trimmed to the region of interest and sectioned the blocks at around 70 nm thickness using a DiATOME diamond knife. The sections were collected onto formvar TEM specimen grids at the different depth as sectioning progressed deeper into the resin block. Thin sections on the TEM grids were stained with 5% uranyl acetate in 50% ethanol for 3 minutes, followed by washing the grids three times with pure water and blotting away the excess water with filter paper. Thin sections on the TEM grid were further stained with 3% Reynold’s lead citrate for 1 minute, followed by washing the grids three times with pure water and blotting away the excess water with filter paper. All grids were air-dried and kept in a grid box for TEM imaging. TEM images were acquired using either a JEOL 1400 flash TEM or HITACHI HT7700 TEM operated at 80 KeV.

### Mass Spectrometry identification of phosphorylated amino acids in HIV particles

Phospho-proteomic analysis of HIV was realised as a fee-for-service at the protein and proteomics core at Children’s Hospital of Philadelphia Research Institute, Philadelphia, Pennsylvania, USA, analogous to previous HIV phospho-proteomic analyses ^16,41^. The mass spectrometry proteomics data have been deposited to the ProteomeXchange Consortium via the MassIVE repository with the dataset identifier MSV000100677. Data are available for review using the following credentials: username: MSV000100677_reviewer, password: 7XqB202rRTuosz33. Details of the phospho-proteomic analyses are as follow:

#### Protein hydrolysis

Purified virus aliquots were removed from storage at -80°C and placed on ice. Lysis buffer (3% SDS, 30 mM Tris pH 7.8, 7.56mM MgCl_2_, 1mM EDTA, 10mM NaF, 1mM PMSF) with protease and phosphatase inhibitors (Sigma P2714, Roche 04 906 837 001) was used to disrupt the concentrated viruses (∼1 mL/50 uL virus pellet) and heated at 90°C for 5 min. The samples were incubated with 5 Units of benzonase (Novagen 70664-3) for 10 min at room temperature. Cysteines were alkylated by the addition of 50 mM iodoacetamide and kept in the dark for 30 min. Proteins were precipitated with 5 volumes of acetone and kept at - 20°C for 2 hours to overnight. Proteins precipitant was centrifuged (14,000 x g, 15 min) and the pellet washed 2x with 80% acetone. The proteins were dissolved in 0.2% SDS, concentration determined by BCA assay, and stored at -80°C. The proteins were digested with 40 µg trypsin (Promega V511A) in 500 uL of 40 mM NH_4_HCO_3_ and 0.1% Rapigest acid labile surfactant (Waters 186001861) at 37°C overnight. Rapigest was hydrolysed with formic acid (2.5% v/v final concentration) for 1 hour at room temperature and centrifuged at 14,000 x g for 20 min. Tryptic peptides were cleaned with a C18 Sep-Pak (WAR036820) eluting with 2 mL 75%,0.1% formic acid. Peptide concentrations were estimated by UV spectrophotometry using an extinction coefficient of 1 mg^-1^cm^-1^. Peptide yields were ∼50% based on the amount of protein input.

#### Hydrophilic interaction chromatography

The protocol of McNulty and Annan for phospho-proteome characterization utilizes hydrophilic interaction chromatography (HILIC) as a first-dimension separation of tryptic peptides, the idea being that the more hydrophilic phospho-peptides are separated from the non-phospho-peptides thus facilitating capture by immobilized metal affinity chromatography (IMAC) with high selectivity ^42^. HILIC was performed on a Beckman-Coulter System Gold HPLC with the following conditions: column, TSK gel Amide 80 4.6 mm x 250 mm (Tosoh Biosciences); Buffers, A-0.1% TFA in water, B-90% CH_3_CN 0.1% TFA; Flow, 0.5 mL/minute; Equilibrate column 85% B 15% A, 1 to 2 mg peptide in 0.5 mL 90% CH_3_CH 0.1%HCOOH loaded at 85% B for 10 min; Gradient, 85% ti 70% B over 40 min, 70% to 10% B over 5 min, hold 10% B 5 min, return to 85% B over 2 min; Collect 2 min fraction from t = 5 to 65 min in 1 mL deep well plate (Eppendorf C5096-0112). Ten percent of each fraction was used for phospho-peptide enrichment.

#### Phospho-peptide Enrichment

Phospho-peptides were enriched from the HILIV fractions using immobilized metal affinity chromatography in batch mode. Phos-Select Iron Affinity beads (Sigma P9740) were added directly to the HILIC fractions (50 uL of 20% evenly suspended slurry) and mixed end over end for 30 min at room temperature. Fractions were transferred to 0.22 µm centrifuge filter devices and centrifuged to remove the filtrate. Beads were washed with 300 µL 30% CH_3_CN, 250 mM AcOH, followed by a wash with water. Filtrates were discarded and phospho-peptides eluted from the beads with 150 µL 400 mM NH_4_OH. After 10 min incubation, filtrates were recovered and lyophilized. Samples were reconstituted with 13 µL 0.15 HCOOH for analysis by mass spectrometry.

#### Mass Spectrometry Analysis

Tryptic digests were analysed on a hybrid LTQ Orbitrap mass spectrometer (Thermofisher Scientific, San Jose, CA) coupled with a NanoLC pump (Eksigent Technologies) and autosampler. Cellular material from one donor was processed (as indicated above) and two sets of MS runs were conducted as technical replicates from both HILIC and IMAC fractions. Tryptic peptides were separated by reverse phase (RP)-HPLC on a nanocapillary column, 75 µm id x 20 cm Protopep (New Objective, Woburn, MA, USA). Mobile phase A consisted of 1% methanol/ 0.1% formic acid and mobile phase B of 1% methanol/ 0.1% formic acid/ 79% acetonitrile. Peptides were eluted into the mass spectrometer at 300 nL/min with each RP-LC run comprising a 15 min sample load at 3% B and a 90 min linear gradient from 5 to 45% B. The mass spectrometer was set to repetitively scan m/z from 300 to 1800 (R = 100,000 for LTA-Orbitrap XL) followed by data-dependent MS/MS scans on the ten most abundant ions, with a minimum signal of 1000, isolation width of 2.0, normalized collision energy of 28, and waveform injection and dynamic exclusion enabled. FTMS full scan AGC target value was 1e6, while MSn AGC was 5e3, respectively. FTMS full scan maximum fill time was 500 ms, while ion trap MSn fill time was 50 ms, microscans were set at one. FT preview mode, charge state screening, and monoisotopic precursor selection were all enabled with rejection of unassigned and 1+ charge states.

#### Sequence database searching and data processing

Raw tandem mass spectrometry data of the phospho-peptide enriched HIV fractions were processed using Proteome Discoverer (version 2.2, Thermo Fisher Scientific) ^43^ with the Sequest HT search engine against the human protein database (version 3.52), and the HIV-1 NL4.3 reference proteome (GenBank accession AF324493.2). Search parameters included trypsin as the proteolytic enzyme allowing up to two missed cleavages, precursor mass tolerance of 10 ppm, and fragment ion mass tolerance of 0.2 Da. Carbamidomethylation of cysteine was specified as a fixed modification, while oxidation of methionine, and phosphorylation of serine, threonine, and tyrosine were set as variable modifications. The Percolator node was used to estimate peptide-spectrum match (PSM) confidence, controlling the false discovery rate (FDR) at 1% at the peptide and protein levels. Phosphorylation site localization was assessed using the IMP-ptmRS node with default settings. Label-free quantitation (LFQ) was performed using precursor ion area detection. The processed data were subsequently reanalysed using Byonic (ProteinMetrics, version 5.4) ^44^ for enhanced peptide-spectrum matching visualization and graphical output generation, which were used in the preparation of figures presented in this manuscript. Similarly, raw MS files from recombinant proteins were processed using Max Quant (version 1.013.13) ^45^. The msm output files were searched against the International Protein Index human protein sequence database (version 3.52, concatenated with reversed decoy sequences and contaminants) and HIV NL4.3 (GenBank AF324493.2) using MASCOT search algorithm (Matrix Science, version 2.3). Fragment ion tolerance was set to 0.7 Da, with a maximum of two missed tryptic cleavage sites. S-Carbamidomethyl cysteine was defined as a fixed modification while oxidized methionine, phospho-serine, phosphor-threonine, and phosphor-tyrosine were selected as variable modifications. The false-discovery rate for peptides and proteins was set at 0.01. The posterior error probability (PEP) threshold was set to 0.05. The re-quantification box in Max Quant was left unchecked to minimise the likelihood of false quantification.

