## Supplementary Data for Bremaud et al for "Calcium-phosphate bridge is a novel phosphorylation switch that stabilises protein-complexes during HIV assembly"

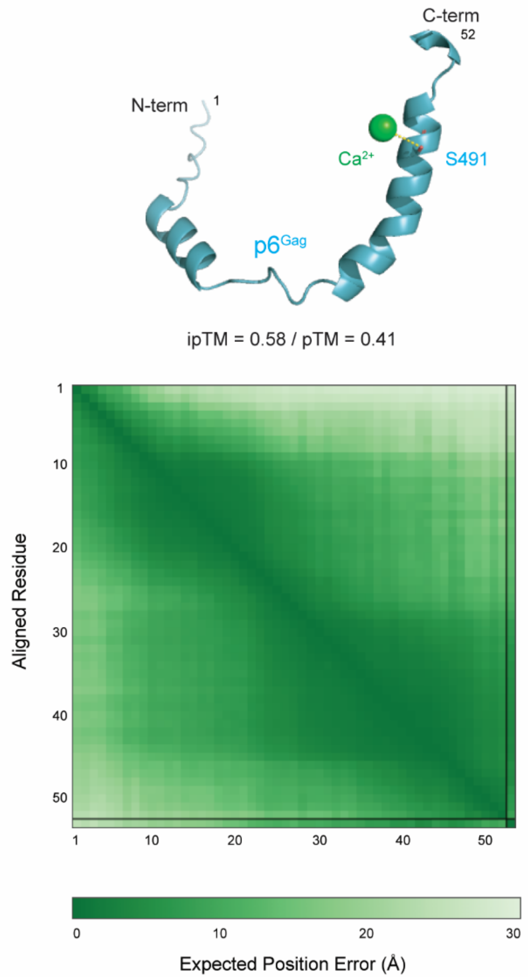


**Supplementary Figure 1: AlphaFold 3 predicted $p6^{Gag}$-$Ca^{2+}$ complex suggests $Ca^{2+}$ likely binds $p6^{Gag}$ near S491 residue.**

AlphaFold server reports that “PAE and/or pLDDT may be more indicative of prediction accuracy [whilst] pTM is less useful for small structures and short chains. This is because the TM score is very strict for smaller molecules”, hence average pLDDT > 70 is presented in Figure 1. Expected position error matrix also highlights high predicted position accuracy, particularly in both predicted alpha-helical domains, whilst ipTM = 0.58 / pTM = 0.41

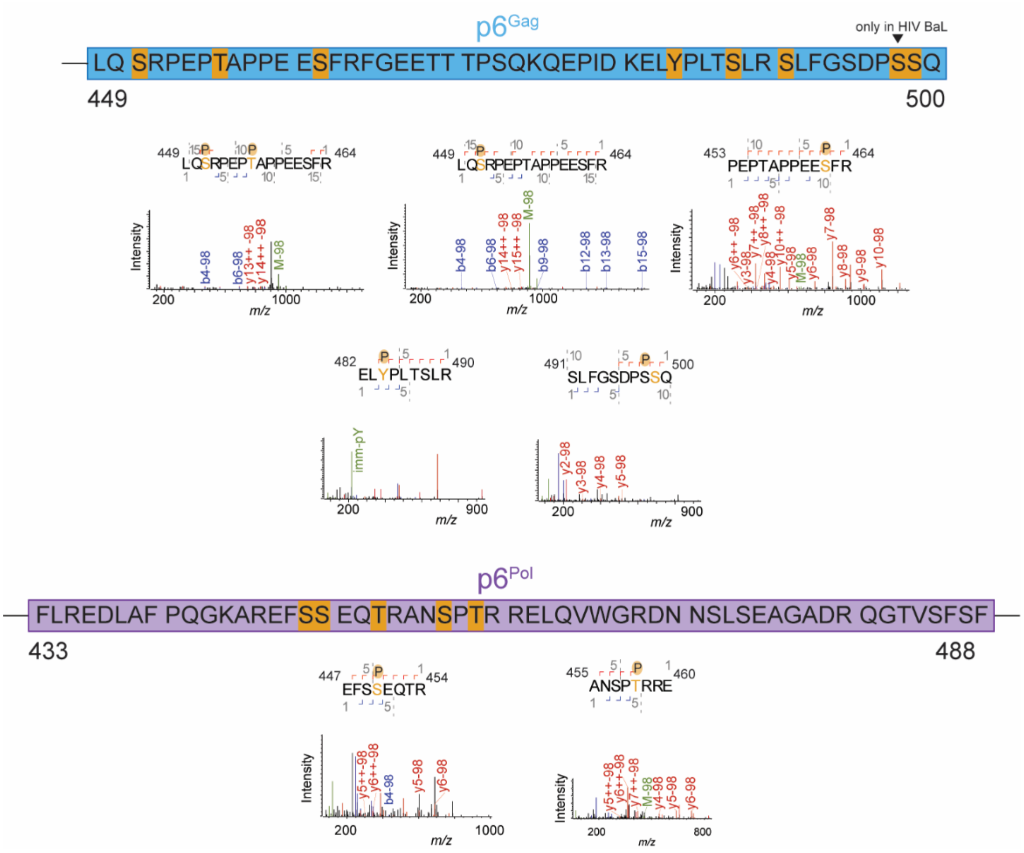


**Supplementary Figure 2: Example of MS spectra displaying evidence of discovered/confirmed sites phosphorylation in p6$^{Gag}$ and p6$^{Pol}$.**

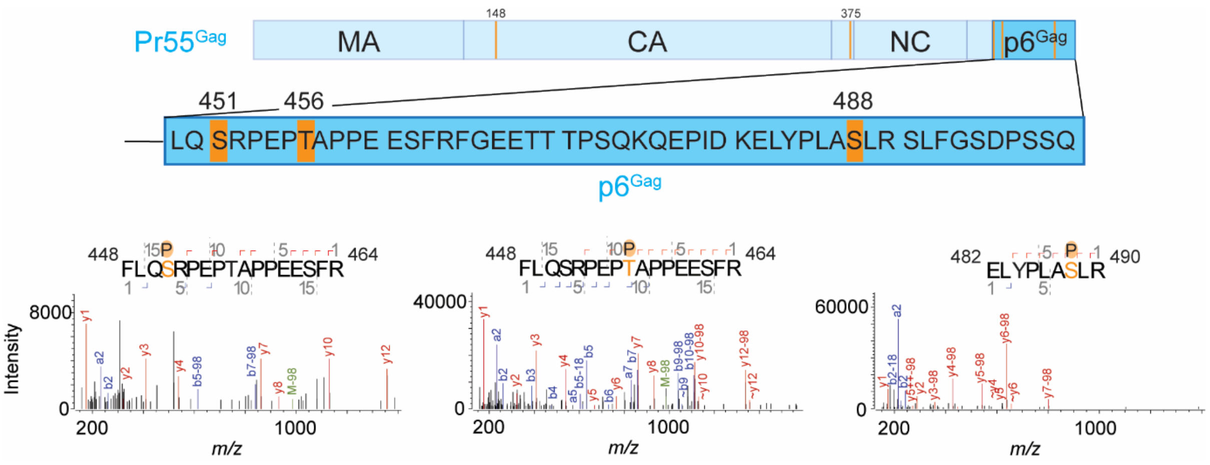


**Supplementary Figure 3: Recombinant Pr55$^{Gag}$ phosphorylation sites and levels upon *in vitro* phosphorylation using active ERK-2.**

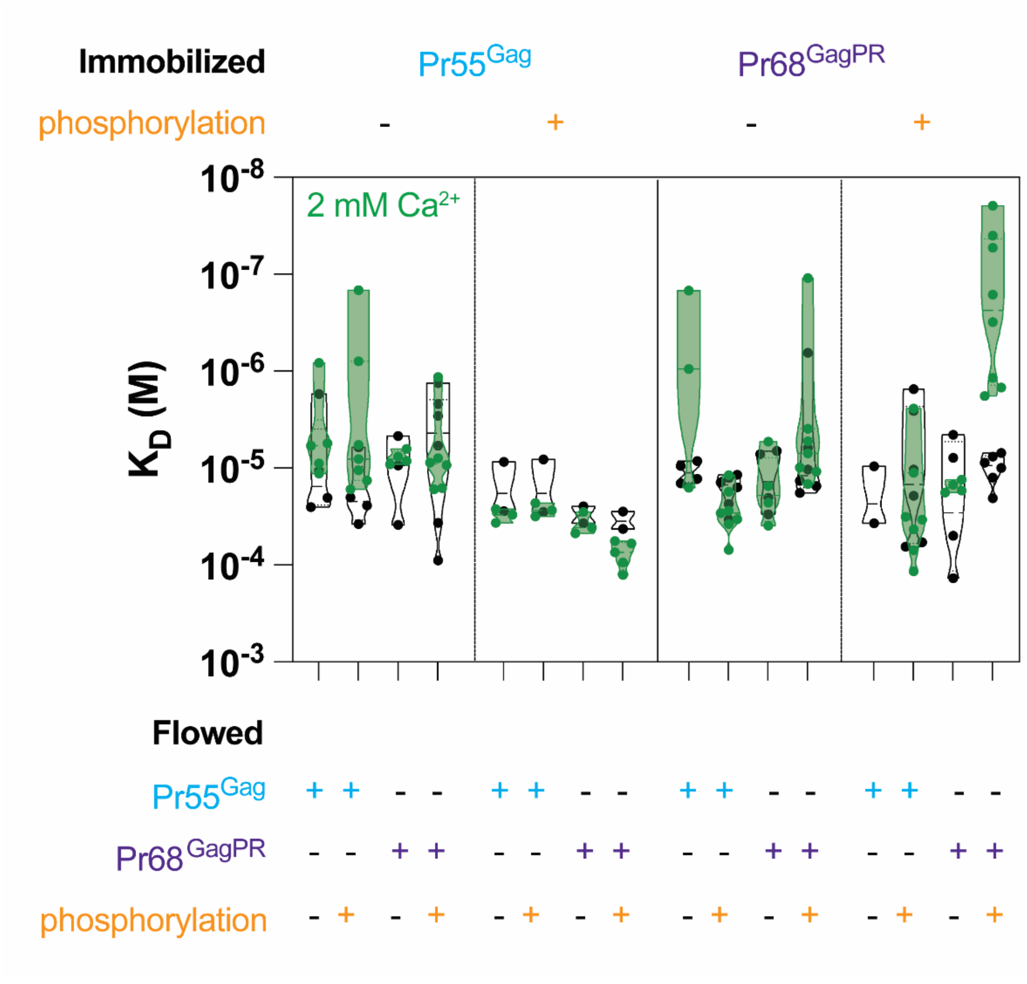


**Supplementary Figure 4: SPR measurements of protein-protein interactions $K_D$ with/without $Ca^{2+}$ mediation.**

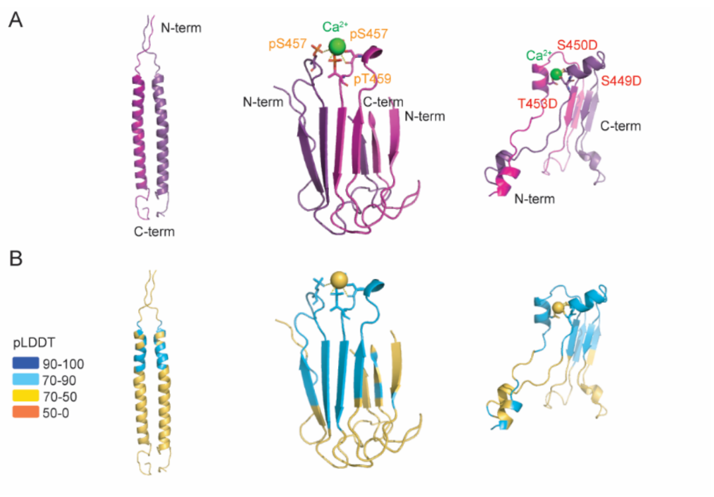


**Supplementary Figure 5: $Ca^{2+}$-phosphate bridges in $p6^{Pol}$ dimer predictions using AlphaFold 3.** (A) Predicted $p6^{Pol}$ dimer structure compared to $Ca^{2+}$ mediated phosphorylated or phospho-mimicking $p6^{Pol}$ dimers. Five sites (S449, S450, T453, S457, T459) are phosphorylated / mutated into D in both $p6^{Pol}$ peptides. (B) Local confidence level of AlphaFold 3 predictions have pLDDT value of > 70 for residues in regions covering S449-T459 in all three simulated dimers composed of 55 residues-long monomers.

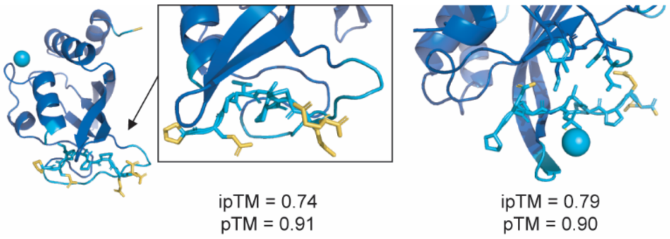


**Supplementary Figure 6: Influence of T456 phosphorylation on Tsg101 UEV domain complex with p6$^{Gag}$ fragment containing PTAP predicted structure in presence of $Ca^{2+}$ using AlphaFold 3.**

Influence of pT456 in PEPTAPPEE p6$^{Gag}$ peptide on predicted complex structure with Tsg101 UEV in presence of $Ca^{2+}$. Proteins structures and interactions predictions are trustworthy (ipTM ≥ 0.74 and pTM ≥ 0.90 in both predictions). Locally, pLDDT ≥ 70 near investigated residues in both predictions.
